# Architecture of the Gβγ-prefusion SNARE complex reveals the molecular mechanism of inhibition of vesicle fusion

**DOI:** 10.64898/2026.03.06.710172

**Authors:** Anna R. Eitel, Montana Young, Jackson Cassada, Eric W. Bell, Jens Meiler, Heidi E. Hamm

## Abstract

Presynaptic inhibitory GPCR (G_i/o_ GPCR) signaling is an essential regulatory mechanism in vertebrate physiology. Near the presynaptic active zone, G_i/o_ GPCR activation releases G-protein βγ heterodimers (Gβγ) which act to inhibit synaptic vesicle fusion through either modulation of Ca^2+^ entry via voltage-gated Ca^2+^ channels, or by direct interactions with the core exocytotic machinery comprised of the ternary SNARE complex downstream of Ca^2+^ influx. The precise molecular mechanism underlying Gβγ-SNARE mediated inhibition has remained unclear due to lack of structural data for the Gβγ-SNARE complex. We address this long-standing question here by stabilizing the interaction between Gβ1γ2 and a pre-fusion ternary SNARE mimetic and determining the structure using single-particle cryo-EM. We used our cryo-EM envelope to build an atomic level prediction of the interaction interface. We validated key interaction sites predicted by our model at the C-terminus of SNAP-25 using site directed mutagenesis and biochemical affinity measurements. Additionally, we found that Gβ1γ2 and a fragment of the regulatory protein complexin can engage the pre-fusion SNARE complex simultaneously. On the basis of these results, we propose a model in which Gβ1γ2 acts on the partially zipped SNARE complex at a late stage in the vesicle docking and priming cycle. In the model, the amino-terminal coiled-coil of Gβ1γ2 forms an interface with the C-terminus of the target membrane SNARE (t-SNARE) complex to prevent complete incorporation of the vesicle SNARE (v-SNARE) into the core SNARE helical bundle, thus blocking vesicle approach to the plasma membrane. The β-propeller domain of Gβ1 may also sterically hinder vesicle approach. Together these results provide crucial structural insights into the mechanism of binding of Gβγ to the SNARE complex, and lends essential insights into the critical role of GPCR signaling to the SNARE complex in modulating synaptic vesicle fusion.

## Introduction

Presynaptic inhibitory G-protein Coupled Receptor (G_i/o_ GPCR) signaling plays an essential role in nervous system function by preventing the release of neurotransmitters.^1^ In response to activation, G_i/o_ GPCRs couple to heterotrimeric G-proteins within the G_i/o_ class. This leads to activation and dissociation of the G-protein into G*α*i, which inhibits adenyl cyclase, and the G-protein βγ heterodimer (Gβγ), which inhibits voltage gated calcium channels as well as exocytotic release machinery.^2^ Inhibition of voltage-gated Ca^2+^ channels suppresses neurotransmitter release at each action potential by limiting Ca^2+^ influx. Gβγ can also bind the SNARE (soluble N-ethyl maleimide sensitive factor attachment protein receptor) complex at a site overlapping the Ca^2+^-synaptotagmin-1 (Syt1) binding interface, thereby inhibiting Ca^2+^-triggered vesicle fusion downstream of Ca^2+^ influx^3-6^.

The latter mechanism of inhibition is transient during trains of stimuli due to residual Ca^2+^ buildup, which restores Syt1 binding and relieves Gβγ-mediated inhibition. We have shown previously that this effect can be reversed by injection of the slow Ca^2+^ chelator EGTA.^7^ As a result, direct inhibition of the SNARE complex by Gβγ provides a rapid and acute mechanism for controlling evoked release while leaving secondary effects of presynaptic Ca^2+^ largely unaffected. Notably, the two Gβγ-mediated inhibitory mechanisms--modulation of Ca^2+^ channels and direct interactions with the SNARE complex--act synergistically during trains of stimuli, enabling complex and dynamic regulation of vesicle fusion associated with Ca^2+^ signaling by distinct GPCRs, with different temporal dimensions to their inhibition.^1,8,9^

Inhibition of neurotransmission via direct interactions between Gβγ and the SNARE complex reduces postsynaptic responses by ∼30-90% depending on the receptor. This mechanism provides a rapid means of tuning synaptic strength and limiting neurotransmitter release during inhibitory GPCR signaling. Removal of this brake on exocytosis in mice results in numerous phenotypes including resistance to diet induced obesity.^4,10,11^ However, the molecular mechanism underlying inhibition of fusion by Gβγ-SNARE interactions remains unclear due to lack of structural information.

The essential machinery required for synaptic vesicle fusion is made up of a family of highly conserved proteins called SNAREs.^12^ The neuronal SNAREs SNAP-25 and syntaxin1a form a binary complex, referred to as target SNARE (or tSNARE), and are anchored to the presynaptic membrane. Synaptobrevin-2 (VAMP2) is anchored to the vesicle membrane. Upon vesicle docking it forms a highly stable ternary complex with SNAP-25 and syntaxin1a referred to as ternary SNARE^13^. The SNARE domains of each protein assemble into a four α-helical bundle in the N-to C-terminal direction to pull membranes toward each other during vesicle docking to the presynaptic membrane (**Fig. 1b**). The docked SNARE complex is primed for fusion through interactions with various regulatory proteins including the Ca^2+^ sensor synaptotagmin-1 (Syt1) and complexin (Cplxn). Upon an action potential, Ca^2+^ interaction with Syt1 and membrane lipids lead to synchronous membrane fusion.^14^

**Figure 1.**
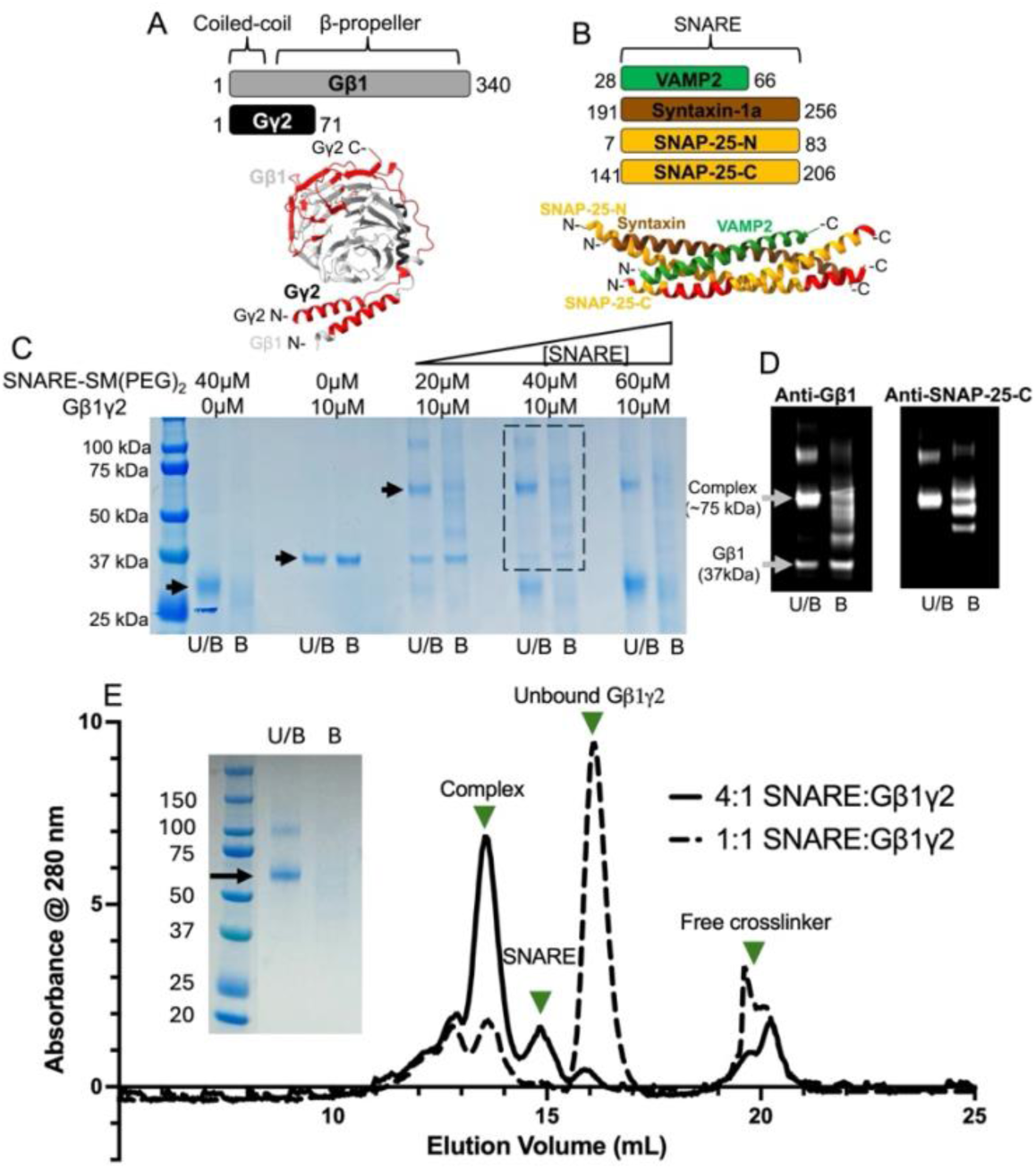
Preparation of stable Gβ1γ2-SNARE complexes using covalent crosslinking. **(a, top)** Domain organization of the Gβ1γ2 heterodimer. (**a, bottom**) Crystal structure of the Gβ1γ2 heterodimer (PDB: 6CRK, from ref. 27). Gβ1 (gray) contains an N-terminal α-helix which forms a coiled-coil with the N-terminus of Gγ2 (black). Gβ1 also contains a β-propeller domain with seven blades. The regions of Gβ1γ2 that bind to prefusion SNARE are highlighted in red (see ref. 17). (**b, top**) Domain organization of the prefusion SNARE complex which contains a C-terminal truncation in the SNARE domain of VAMP2 (green), mimicking a partially zipped conformation. (**b, bottom**) Crystal structure of the pre-fusion ternary SNARE complex (PDB: 5W5D, from ref 23). Syntaxin-1a is shown in brown and the SNAP-25 N-terminal (SNAP-25-N) and C-terminal (SNAP-25-C) helices are shown in orange. The regions of SNAP-25 that bind Gβγ determined via peptide array analysis (see ref. 28) are shown in red. Note that only a small portion of the essential extreme C-terminal binding site of SNAP-25-C is resolved in the crystal structure likely due to the lack of order in this region. **(c)** SDS-PAGE analysis of crosslinking conditions. SM(PEG)₂-SNARE to 10 µM Gβ1γ2 ratios ranged from 2:1 to 6:1. “U/B” samples were unboiled to assess SNARE assembly; the intact ternary SNARE band (∼29 kDa) appears only in U/B samples. The Gβ1 band (37 kDa) decreases with higher SNARE concentration as it is consumed in crosslinking. The dashed box indicates the region shown in (d). **(d)** PVDF membranes probed for Gβ1 (left) or SNAP25 (right). Gβ1 and SNAP25 co-migrate at ∼75kDa (gray arrow); a faint ∼120kDa band may represent the complex at a greater stoichiometry. **(e)** Size-exclusion profiles of SM(PEG)2-crosslinked Gβ1γ2-SNARE on a Superdex 200 Increase 10/300 GL column. Samples contained 100µL of 10µM Gβ1γ2 with 10µM (solid black) or 40µM (dashed line) SM(PEG)2-SNARE. The free Gβ1γ2 peak (∼16mL) decreases and the Gβγ-SNARE peak (∼13.5mL) increases with excess SNARE. (inset) SDS-PAGE of the complex fraction used for cryo-EM.

Gβγ interacts with all 3 neuronal SNARE proteins: SNAP-25, syntaxin, and VAMP2.^3^ The C-terminus of SNAP-25 is the most important interaction site and is essential for Gβγ-SNARE mediated inhibition in vivo.^3,5^ We have shown that Gβγ competes with the Ca^2+^ sensor Syt1 for binding to the SNARE complex,^3,6,15^ however, it may also directly block access of the vesicle to the plasma membrane (PM) sterically due to its binding to the C-terminal end of the SNARE complex, the site of vesicle approach to the membrane. Interference with Gβγ-SNARE interactions with a peptide mimicking the 14 c-terminal residues of SNAP-25 does not alter synaptic transmission but prevents Gβγ-mediated presynaptic inhibition.^5^ We also showed that Gβγ can inhibit membrane fusion by inhibiting Syt1 and Ca^2+^-induced lipid mixing in a dose-dependent manner in a reconstituted liposome fusion assay, proving that this is a direct protein-protein interaction.^6^ We have concluded that this region of the SNARE complex is an important site of neuromodulation. Furthermore, the identity of the Gγ isoform largely governs the affinity of Gβγ for SNARE. For instance, Gβ1γ2 has a significantly higher affinity for SNARE than Gβ1γ1,^6,16^ and the peptide corresponding to the N-terminal α-helix of the Gγ2 subunit disrupts the interaction competitively.^17^

There are five Gβ and twelve Gγ isoforms in humans.^18^ Gβ is classified as a WD-40 repeat protein and contains a 7-bladed β-propeller domain and an N-terminal α-helix that forms a coiled-coil with the N-terminus of Gγ.^19,20^ The primary effector interaction site of βγ involves the top of the β-propeller domain of Gβ, which is also where switch 2 of Gα interacts in the inactive Gα(GDP)βγ heterotrimer.^21^ Gγ is comprised of two α-helices which interact with the Gβ subunit and a disordered C-terminus which is prenylated to target Gβγ to the membrane. Our previous peptide array analysis identified binding sites for both the t-SNARE (syntaxin/SNAP-25) and ternary SNARE complex on Gβ1γ2, which include both the N-terminal coiled-coil and β-propeller domains of Gβγ (**Fig. 1a**).^17^

Despite extensive characterization of this mechanism of inhibition through both biochemical and *in vivo* studies, the molecular mechanism of Gβγ-SNARE mediated inhibition remains unclear due to lack of structural data. Additionally, the consequences of Gβγ binding on interactions between the SNARE complex and other binding partners, such as complexin and Munc18, remain undetermined. To address this issue, we stabilized the interaction between soluble Gβ1γ2 and a partially zipped pre-fusion ternary SNARE mimetic and determined the structure of the intact complex using single-particle cryo-EM. Based on our preliminary envelope, we generated a computational model of the interface and used it to generate SNAP-25 C-terminal mutants with increased affinity for Gβ1γ2. Our computational model based on our structural envelope is in good agreement with our previously published biochemical studies and reveals that the N-terminal coiled-coil of Gβγ forms an interface with the C-terminus of the SNARE complex, through key residues of SNAP-25. The overall architecture of the complex reveals that Gβγ acts at the C-terminus of the t-SNARE complex during vesicle docking to disrupt SNARE complex zippering and would sterically hinder vesicle approach and full collapse of the vesicle membrane.

## Materials and Methods

### Protein purification

His-tagged soluble Gβ1γ2 (C68S) was expressed and purified from Sf9 cells essentially previously described,^22^ however, the Gβ1γ2 construct contained a hexahistidine tag on the N-terminus of Gβ1. Except for Gβ1γ2 used for MST and Alphascreen assays, the His tag was cleaved with the TEV protease prior to anion exchange chromatography. For the soluble ternary SNARE complex, plasmids encoding rat 6xHis-tagged synaptobrevin-2 (residues 28-66) and syntaxin-1A (residues 191-256) in a pACYDuet-1 were received as gifts from the laboratory of Dr. Qiangjun Zhou (Addgene plasmid #70055, #106414). Plasmids encoding rat SNAP-25b (residues 141-206 and 7-83) were subcloned into a pETDuet-1 by GenScript. SNARE-containing plasmids were co-transformed into BL21 *E. Coli*. Purification of the SNARE complex was performed as previously described.^23^ Elution from the anion exchange column (Cytiva HiTrap-Q, 5 mL) occurred in two peaks, the first peak that elutes at a conductivity around 20 mS/cm contains the properly folded ternary SNARE complex. All mutant ternary SNARE complexes were purified exactly as the wildtype. The plasmid encoding GST-tagged rat complexin (residues 1-83) was purchased from Addgene (plasmid #106417),^23^ expression and purification was performed as previously described.^23^

### Microscale thermophoresis

For MST experiments, the N-termini of Gβ1γ2 was fluorescently labeled with Alexa Fluor 647 NHS ester (AF647) using a protocol which selectively targets amino termini of proteins.^24^ The degree of labeling was estimated based on the absorbance values at 650 and 280 nm. Typical values were 1.8 molecules of AF647 per molecule of Gβ1γ2. AF647-conjugated Gβ1γ2 was diluted to a protein concentration of 10 nM in assay buffer (PBS + 0.05% Tween-20). The wildtype and mutant partially zipped ternary SNARE mimetics were diluted to a final concentration of 60 μM and serially diluted in PCR tubes with 10 μL of assay buffer. Following centrifugation at 13,000 x g, 10 nM AF647-Gβ1γ2 was added to each PCR tube. The final volume in each tube was 20 μL. Samples were immediately placed in Nanotemper standard capillary tubes and read three times on a Nanotemper Monolith Pico MST instrument. For each condition, samples were prepared fresh three times and read three times on the MST instrument to give a total of nine measurements. For the complexin 1-83 experiments, the prefusion SNARE complex was pre-incubated with equimolar concentrations of complexin 1-83 prior to MST analysis.

### Chemical crosslinking

The SM(PEG)_2_ crosslinker (Pierce, 1 mg no-weigh format, ThermoFisher catalogue # A35397) was dissolved in DMSO to a final concentration of 25 mM. Partially zipped SNARE was exchanged into conjugation buffer (PBS, pH = 7.3, 1 mM EDTA) prior to addition of the stock SM(PEG)_2_ solution to a final concentration of 1 mM. After 30 minutes, unreacted crosslinker was removed via desalting column (Zeba spin desalting column, 500 μL, 7K MWCO). Various concentrations of SM(PEG)_2_-conjugated SNARE were then incubated with 10 μM soluble Gβ1γ2 overnight. The following day, samples were assessed via SDS PAGE and gel filtration chromatography using a Superdex 200 Increase 10/300 GL column equilibrated with conjugation buffer.

### Cryo-electron microscopy

SEC fractions corresponding to the crosslinked Gβ1γ2 SNARE complex were pooled and concentrated to 0.17 mg/mL and 2.5 μL of sample was immediately applied to UltrAuFoil grids (Quantifoil, R 1.2/1.3) at 7°C and 100% humidity using a Vitrobot IV. Optimal freezing conditions were blot time = 10.5 seconds and blot force = 12. ∼30,000 cryo-EM movies were acquired on a Titan Krios 300 kV cryo-TEM with a Gatan K3 direct electron detector using the following parameters: magnification: 215,000x, unbinned pixel size: 0.57886 -Å, spherical aberration: 2.7 mm, spot size: 4, parallel beam size: 500 nm, dose rate: 7.573 e-/pixel/s, exposure time: 2.651 seconds, total dose: ∼60 e-/Å2, 846 frames in EER format, defocus range: -2.2 to 0.9 μm in 0.1 μm steps.

### Cryo-EM data processing

Movies were imported into cryoSPARC. Preprocessing was performed using patch motion correction and patch CTF estimation. Peak SEC fractions corresponding to the complex were immediately vitrified. After systematic screening of grid type, and blotting force and time, we identified conditions that reproducibly produced quality vitrified samples of the crosslinked Gβ1γ2-SNARE complex. The best-quality grid was submitted for data acquisition on a ThermoFisher Krios 300 kV cryo-TEM with a Gatan K3 direct electron detector. Approximately ∼32,000 movies were acquired and processed with cryoSPARC. Following motion correction and CTF estimation, micrographs with a CTF fit from 2.5-10-Å, defocus range of >20, and an average intensity of >5.4 were accepted, resulting in a final count of 22,000 micrographs to be used for particle picking. ∼1000 particles were picked manually and used to train the TOPAZ machine learning particle picking program. This model was used to pick ∼1 million particles from the entire dataset.

## Results

### Preparation of stable Gβ1γ2-SNARE complexes for structure determination

Soluble ternary SNARE mimetics were used for structural studies as they are stable and have been previously crystallized successfully.^14,23,25^ The partially zipped ternary SNARE mimetic^23^ was previously shown to bind to Gβ1γ2 with a higher affinity than that of the fully zipped mimetic, ^17^therefore, this construct was used for our structural studies. This construct contains only the SNARE domains of rat syntaxin-1a (residues 191-256, brown) and SNAP-25 (residues 7-83 and 141-206, orange) and contains a portion of the VAMP2 (synaptobrevin-2) SNARE domain (residues 28-66 green, **Fig. 1b**). The VAMP2 SNARE domain contains a C-terminal truncation which prevents SNARE complex zippering and mimics the partially zipped, pre-fusion state.^23^

Our attempts to determine the Gβγ-SNARE complex structure via cryo-EM without crosslinking have been unsuccessful. The particles did not align into 2D classes, indicating that the Gβγ and SNARE were flexible with respect to one another. To increase rigidity at the interaction interface, we crosslinked the Gβγ-SNARE complex with SM(PEG)_2_ **(Fig. 1c-e)**. This is a heterobifunctional crosslinker reactive towards Lys and Cys residues with a spacer arm length of ∼17-Å. To test crosslinking efficiency, the SNARE complex (that contains no Cys) was reacted with a 10-fold molar excess of SM(PEG)_2_ for 30 minutes at room temperature in PBS, pH = 7.3. Unreacted crosslinker was removed with a desalting column. SM(PEG)_2_-conjugated SNARE complex ranging in concentrations from 20 to 60 μM was incubated with 10 μM Gβ1γ2 overnight under non-reducing conditions. Gβ1γ2 contains 15 Cys residues; the majority are buried within the protein core except for Gβ1 Cys 204. Samples were analyzed via SDS-PAGE (**Fig. 1c**) and western blot probing for Gβ1 and C-terminal SNAP25 (**Fig. 1d**).

The extent of the crosslinking reaction can be assessed by monitoring the intensities of the SDS-PAGE gel bands corresponding to Gβ1 (37 kDa, dark green arrow) and the Gβγ-SNARE complex (∼75 kDa, red arrow) with increasing SNARE concentration (**Fig. 1c**). In SDS sample buffer ternary SNARE complex is resistant to SDS-denaturation in the absence of boiling (**Fig 1c**, bottom)^26^. Samples are labeled boiled (“B”) or un-boiled (“U/B”). The band around 29 kDa in the first “U/B” lane (purple arrow) corresponds to properly folded SM(PEG)2-ternary SNARE. The condition with a 4:1 molar ratio of SM(PEG)2-SNARE complex to Gβ1γ2 is optimal because it contains minimal unbound components (**Fig. 1c**, outlined in box).

We analyzed crosslinked samples containing different molar ratios of SM(PEG)2-conjugated SNARE complex to Gβ1γ2 using size-exclusion chromatography (SEC, **Fig. 1e**). The elution volumes corresponding to Gβ1γ2, SM(PEG2)-SNARE complex, and the crosslinked complex are ∼16 mL, 15 mL, and 13.5 mL, respectively. The most prominent peak in the SEC profile of the 1:1 sample corresponds to free Gβ1γ2 (**Fig. 1e**, blue trace). When the ratio of SM(PEG)2-SNARE complex to Gβ1γ2 was increased to 4:1, nearly all free Gβ1γ2 was crosslinked to the SNARE complex as indicated by the prominence of the complex peak (**Fig. 1e**, red trace).

To ensure the crosslinked samples contained the properly folded ternary SNARE complex and both Gβ1 and Gγ2, we analyzed the SDS-PAGE gel bands of interest with mass spectrometry The indicated bands in Fig 1c were excised and subjected to in-gel trypsin digestion. Resulting tryptic peptides were analyzed via high-resolution LC-MS/MS using the Proteomics Lab of the Mass Spectrometry Research Center at Vanderbilt University. Proteins were identified using Sequest the normalized total spectra corresponding to each protein was determined with Scaffold 5. Indeed, all six protein components were present in the crosslinked complex.

### Architecture of the Gβγ-SNARE complex revealed via cryo-EM

Peak SEC fractions corresponding to the complex were immediately vitrified and submitted for acquisition on a ThermoFisher Krios 300 kV cryo-TEM with a Gatan K3 direct electron detector. Data were processed in cryoSPARC. The final number of micrographs used for particle picking with the TOPAZ particle picking algorithm was ∼22,000. The representative 2D classes of ∼500K curated particles shows a ternary SNARE complex, which appears rod-shaped, bound to Gβγ, which appears globular (**Fig. 2a**). These particles were then sorted into 3D ab-initio classes (**Fig. 2b**) and heterogeneously refined (**Fig. 2c**). The largest class showed density consistent with both Gβ1γ2 and an elongated helical SNARE bundle (**Fig. 2b, c**, teal surface). Additionally, there is a class showing density for Gβ1γ2 alone, with the N-terminal coiled-coil and β-propeller domains clearly distinguished (**Fig. 2b, c**, gray surface). The "Gβ1γ2 only" map was fit into the largest class in ChimeraX resulting in a correlation of 0.88, indicating that our assignment of Gβ1γ2 to the globular density is accurate (**Fig. 2b**, far right).

**Figure 2.**
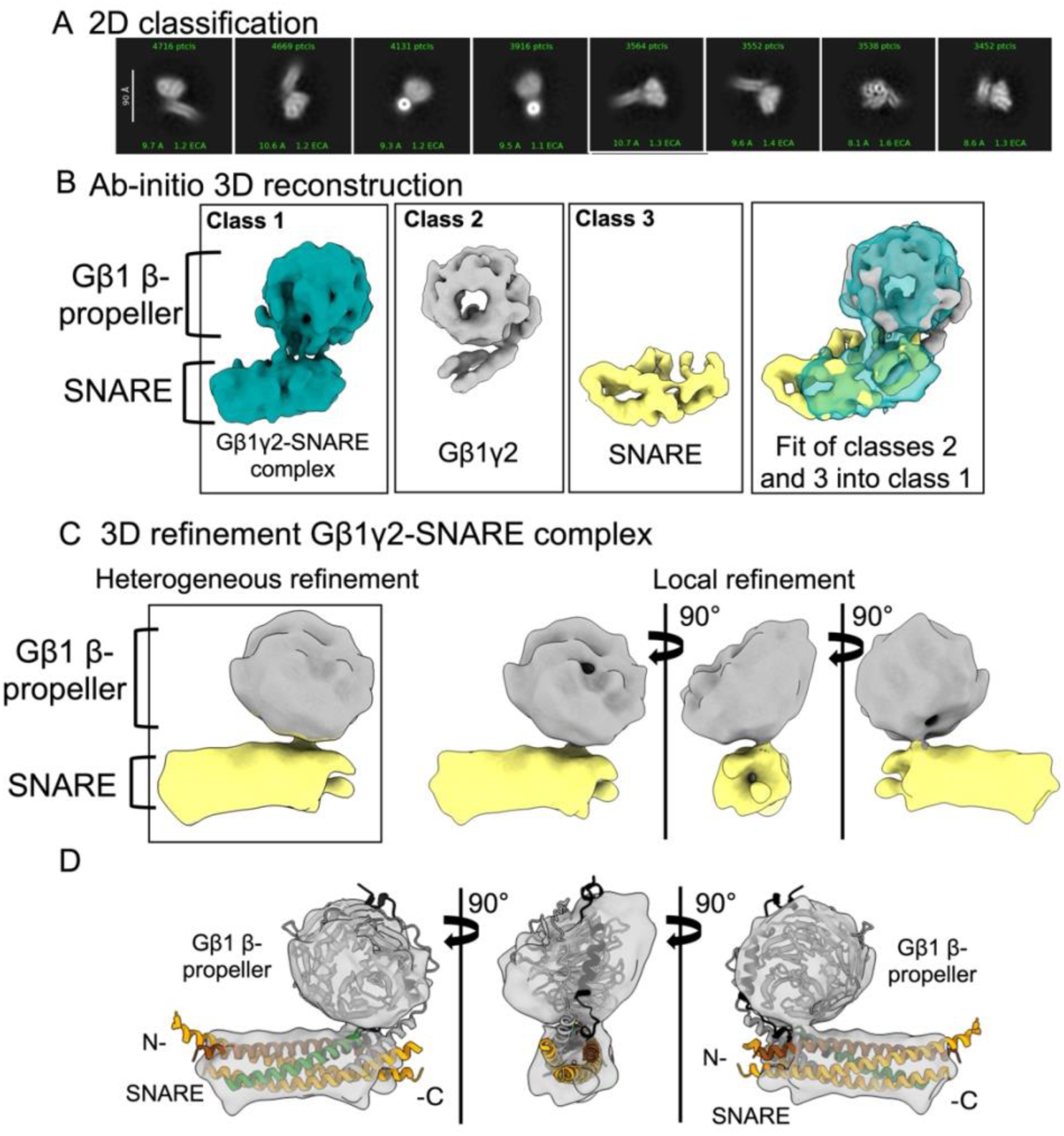
Single-particle cryo-EM of the Gβ1γ2-prefusion SNARE complex. **(a)** Representative 2D classes of ∼500K particle picked via TOPAZ. The pixel size used for import was 0.58-Å. The extraction box size used was 360 pix (∼209-Å). The prefusion SNARE complex appears as a rod-like extension from a circular pattern of density which corresponds to the Gβ1 β-propeller. **(b)** Representative ab-initio 3D classes of the Gβ1γ2-prefusion SNARE complex. The largest class, shown as a teal surface, contains 147K particles and shows density consistent with Gβ1γ2 and an associated SNARE complex. We also observed a class corresponding to Gβ1γ2 alone (gray surface, 86K particles), in which the N-terminal coiled-coil and β-propeller domains are well defined, and SNARE alone (yellow surface, 64K particles). Fitting of the "Gβ1γ2 alone" (class 2) and "SNARE alone" (class 3) maps into the "Gβ1γ2-SNARE complex" (class 1) map (far right) resulted in good correlation. **(c)** 3D refinement of the Gβ1γ2-SNARE complex class. **(d)** Fitting of the Gβ1γ2 (PDB: 6CRK) and pre-fusion SNARE (PDB: 5W5C) crystal structures into the map shown in (c).

Volumes containing density for both Gβ1γ2 and SNARE were then refined via non-uniform 3D refinement (**Fig. 2d**). To increase the resolution, the density was masked around the SNARE and the Gβγ β-propeller for local refinement. We estimate the resolution of our final map to be ∼8-Å based on both the GSFSC curves (**Fig. 2d**).

The crystal structures for Gβ1γ2 (PDB: 6CRK)^27^ and pre-fusion SNARE (PDB: 5W5C)^23^ were fit into the cryo-EM map using rigid body fitting in ChimeraX to determine the overall architecture of the complex within the envelope (**Fig. 2d**). Both crystal structures are in good agreement with the density. Density for Gβγ appears at one end of the SNARE complex, which we hypothesize is the C-terminus, based on the well-established importance of this region for the Gβγ-SNARE interaction and Gβγ-SNARE mediated inhibition of fusion. Our previous studies identified regions of the Gβ1 β-propeller domain, near the main binding site for Gα(GDP), and regions of the N-terminal coiled-coiled of Gβ1γ2 that interact with the SNARE complex (**Fig. 1a, bottom**). Because N-terminal coiled-coil of Gβ1γ2 is not observed elsewhere near the Gβ1 propeller domain, as would be expected based on previous maps containing Gβ1γ2 at this resolution, we hypothesize that the N-termini are buried in the interface with the C-terminal end of the SNARE complex.

### Computational modeling and prediction of high-affinity mutations at the Gβγ-SNARE interface

We next sought to build an atomic resolution computational prediction of the Gβγ-SNARE complex. Structural information from our cryo-EM envelope together with data from our previous studies in which we identified critical sites of interaction,^15,17,28^ has allowed us to generate a molecular model of N-terminal Gβ1γ2 bound to the pre-fusion ternary SNARE mimetic using computational means (**Fig. 3**). Chai-1 and AlphaFold were used to model the complex to maintain the coiled-coil interaction between Gβγ and the prefusion-SNARE complex. Rosetta cartesian relaxation was used to ensure biological feasibility of the generated model^29-34^. Consistent with the cryo-EM envelope and biochemical studies, the Gβ1γ2 N-terminal coiled-coil is predicted to dock at the extreme C-terminus of the SNARE complex and insert into the SNARE helical bundle (**Fig. 3a**).

**Figure 3.**
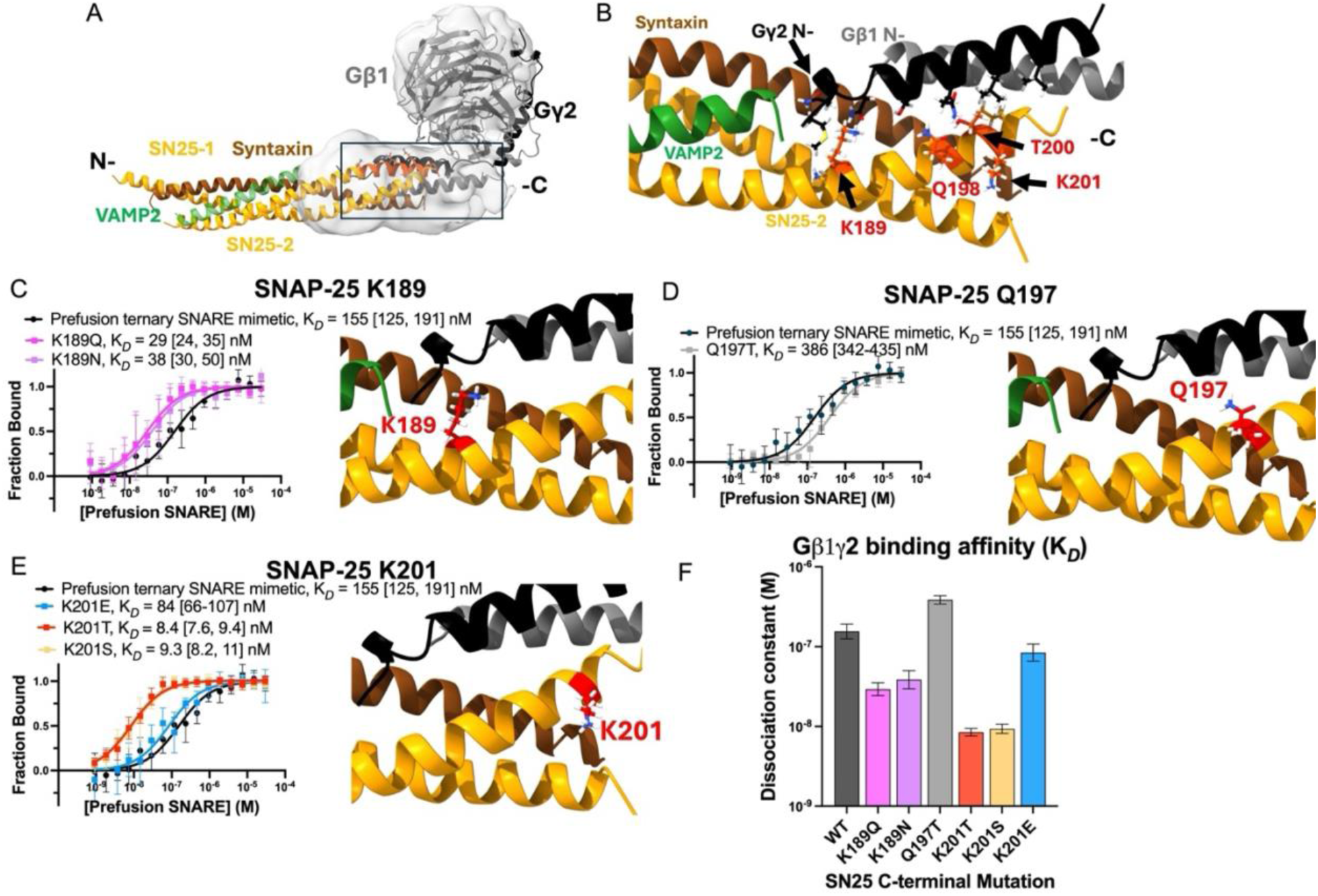
Atomic model of the Gβ1γ2-prefusion SNARE complex and experimental validation. **(a)** Computational model of the Gβ1γ2 N-terminal peptides bound to the pre-fusion SNARE complex. The crystal structure for Gβ1γ2 is superimposed into the model. The envelope from cryo-EM is shown as a transparent gray surface and is in good agreement with the computational model**. (b)** Predicted interface between N-terminal Gβ1 (gray) Gγ2 (black) and the pre-fusion SNARE complex. Positions that were mutated are shown as red sticks. **(c-e)** Microscale thermophoresis analysis of the pre-fusion SNARE complex containing predicted high-affinity mutations binding to 5 nM fluorescently labeled Gβ1γ2. Substitution of SNAP-25 K201 for either a threonine (red) or serine (orange) significantly increased the affinity for Gβ1γ2, as predicted by the model. **(f)** Dissociation constants for each ternary SNARE mimetic calculated from the curves shown in (c-e) using GraphPad Prism.

To probe the residues present at the binding interface, we used this computational model to predict point mutations in the SNARE complex that would increase its affinity for Gβ1γ2. Eight such mutants were predicted at four different positions within the C-terminal helix of SNAP-25 (**Fig 3b)**. The sites targeted for mutagenesis by Rosetta were K189, Q198, T200, and K201 (**Fig. 3b**, red sticks). We expressed and purified these pre-fusion SNARE mutants and compared their Gβ1γ2 binding to wild type using MST (**Fig. 3c-e**). The affinity of the wildtype SNARE to 5 nM fluorescently labeled Gβ1γ2 was determined to be 155 nM (95% CI: 125-191 nM, black circles). We identified two single point mutants with significantly increased affinity for Gβ1γ2, the SNAP25 K201T and SNAP25 K201S mutants, with K*_D_* values of 8.4 nM (95% CI: 7.6-9.4 nM, red squares) and 9.3 nM (95% CI: 8.2-11 nM, orange squares), respectively (**Fig. 3e, f**). The SNAP-25 K189N (purple squares) and K189Q (pink squares) mutants also had an increased affinity for Gβ1γ2 (**Fig. 3c, f)**. The SNAP25 K201E (K*_D_* = 84.3 nM, 95%CI: 65.9-107 nM) and Q197T (K*_D_* = 386 nM, 95%CI: 342-435 nM) showed modest changes in affinity **(Fig. 3, d-f**).

### Complexin is compatible with the Gβγ-SNARE complex

To investigate possible cooperation between Gβ1γ2 and other regulators of fusion, as well as add to add size and stability to the Gβ1γ2-SNARE complex for structure determination, we next assessed the ability of partially zipped SNARE to interact with Gβ1γ2 and complexin simultaneously (**Fig. 4**). Complexin is a small soluble protein that cooperates with SNAREs and synaptotagmin to trigger fusion upon Ca^2+^ influx.^35^ It is known that synaptotagmin and complexin can bind the SNARE complex simultaneously but synaptotagmin and Gβ1γ2 cannot.^6,23^ Therefore, we hypothesized that Gβ1γ2 and complexin simultaneously interact with the SNARE complex. Complexin is known to stabilize the *trans*-SNARE complex^36,37^ to which Gβ1γ2 has increased affinity, therefore we also hypothesized that the presence of complexin would increase the affinity of Gβ1γ2 for partially zippered SNARE.

**Figure 4.**
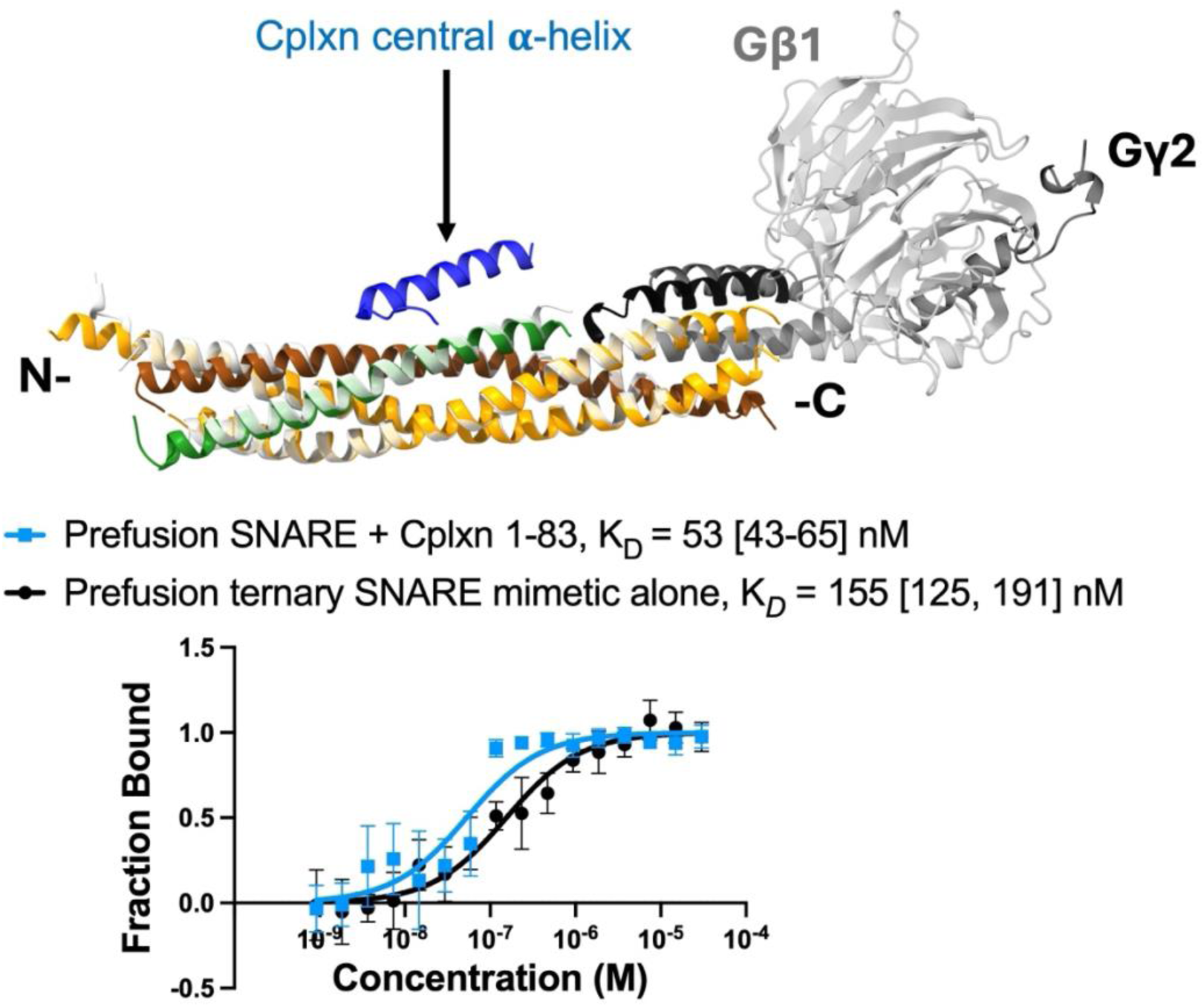
Gβ1γ2 and the Complexin-1 central α-helix (residues 1-83) can engage the pre-fusion SNARE complex simultaneously. **(a)** Crystal structure of the pre-fusion SNARE-complexin 1-83-synaptotagmin complex (PDB: 5W5C, from ref. 23) with our computational prediction superimposed. Synaptotagmin is removed for clarity. The binding site for the central α-helix of complexin (blue) is not expected to overlap with that of the Gβ1γ2 N-termini. **(b)** MST binding analysis of 5 nM fluorescently labeled Gβ1γ2 binding to pre-fusion SNARE in the presence of equimolar concentrations of complexin 1-83 (blue squares). The Gβ1γ2-SNARE binding affinity is slightly higher in the presence of the complexin fragment. The K_D_ in the absence of complexin was determined to be 155 nM (black) and 53 nM in the presence of complexin (blue). This suggests that the binding sites for complexin and Gβ1γ2 on ternary SNARE do not overlap.

We prepared a complexin fragment (res. 1-83) from *E. Coli.* This fragment contains the N-terminal, accessory, and central alpha helical domains, but not the C-terminal domain which interacts with membrane lipids.^23^ The central alpha helical domain binds to the SNARE complex and is essential for all complexin functions (**Fig. 4a**).^38^ Prefusion SNARE and complexin 1-83 were incubated at equimolar concentrations prior to preparation of a concentration response curve. The K*_D_* corresponding to the Gβ1γ2-SNARE-Cplxn complex is 53 nM (95% CI: 43-65 nM) (**Fig. 4b**, blue squares) whereas in the absence of complexin the K*_D_* = 155 (95% CI: 125-191 nM) (**Fig. 4b**, black circles). Therefore, the presence of complexin does indeed appear to increase the affinity of Gβ1γ2 for the pre-fusion SNARE complex. These results provide further mechanistic detail into Gβγ-SNARE mediated fusion by indicating Gβ1γ2 and complexin can bind to the SNARE complex simultaneously. Furthermore, the fact that the complexin 1-83 binding site and the predicted Gβ1γ2 binding site on pre-fusion SNARE do not overlap provides further support for our model (**Fig. 4a**).

## Discussion

Gβγ-SNARE mediated inhibition of synaptic vesicle fusion is an important regulatory mechanism of neurotransmission.^1,3,5,16,39,40^ While this interaction has been studied extensively both biochemically^3,6,15,28,41^ and *in-vivo*,^4,10,11^ the molecular mechanism underlying inhibition remains poorly understood due to the lack of structural data for the complex. Here, we provide the first structural model of the Gβγ-SNARE complex which provides essential insights into the mechanism of inhibition. Our single particle cryo-EM envelope revealed the overall architecture of the complex and indicates that the N-terminal coiled-coil of Gβ1γ2 forms an interface with the C-terminus of the partially zippered SNARE helical bundle. This envelope allowed us to build a high-resolution, testable computational prediction, which we verified experimentally by correctly predicting point mutations at the C-terminus of SNAP-25 that increase the affinity of the pre-fusion SNARE complex for Gβ1γ2. Furthermore, we discovered that partially zipped SNARE can interact with Gβ1γ2 and complexin simultaneously. Taken together, our results suggest a mechanism of inhibition in which the N-termini of Gβ1γ2 insert into the core SNARE helical bundle at the C-terminus of SNAP-25 at the pre-fusion stage and interfere with SNARE complex zippering. We propose that Gβγ acts by completing the helical bundle and sterically hindering the final zippering up of synaptobrevin and vesicle approach to the plasma membrane (**Fig. 5**).

**Figure 5.**
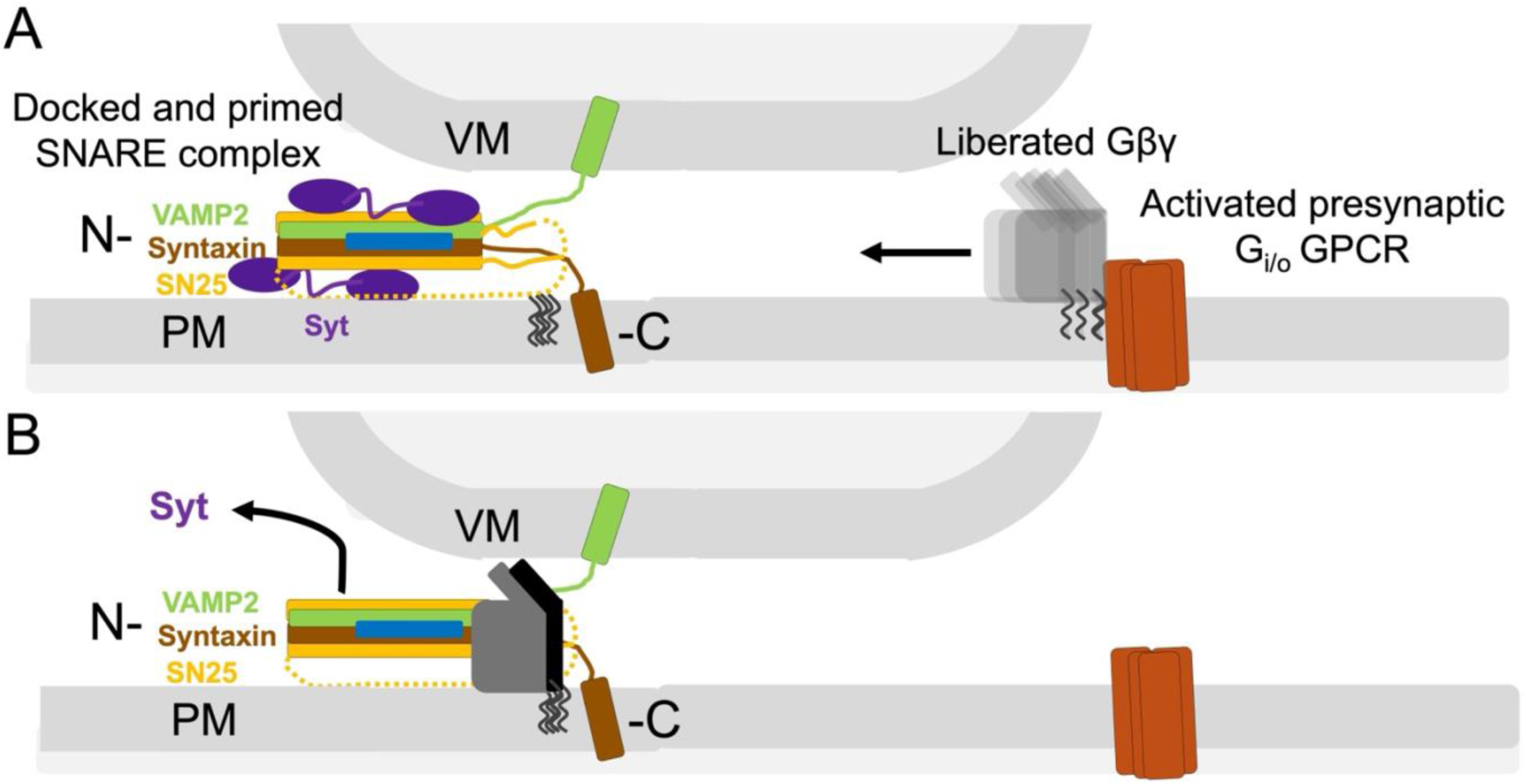
Proposed molecular mechanism of Gβ1γ2-SNARE-mediated inhibition of synaptic vesicle fusion. **(a)** Upon activation of presynaptic Gi/o GPCRs, liberated Gβγ diffuses across the membrane and interacts with the pre-fusion SNARE complex. **(b)** At low Ca^2+^ concentrations, Gβγ displaces synaptotagmin (purple) and inserts into the C-terminus of the t-SNARE helical bundle, thereby disrupting SNARE complex zippering via the Gβγ N-terminal coiled-coil domain. The β-propeller domain may also sterically block vesicle approach.

After an action potential invades the synapse, Gβγ liberated from either autoreceptors that were activated by the first action potential, or neuromodulators activating heteroreceptors, Gβγ binding would stabilize a partially zipped SNARE intermediate and suppress vesicle fusion in conditions of low Ca^2+^. During stimulation, however, rising residual Ca^2+^ concentrations would promote displacement of Gβγ by Ca^2+^-Syt1, driving completion of SNARE complex zippering. This dynamic competition provides a structural explanation for physiological observations that Gβγ-mediated inhibition is relieved during trains of stimuli as intracellular Ca^2+^ accumulates.^7^ This mechanism provides a structural basis for the rapid and reversable inhibition of neurotransmitter release observed during presynaptic G_i/o_-coupled GPCR signaling. Because Gβγ can be rapidly released following GPCR activation^5^ and directly engage the SNARE complex, this interaction provides a means of modulating vesicle fusion independently of presynaptic Ca^2+^ entry. Thus, Gβγ-SNARE interactions represent a critical point of convergence between GPCR signaling pathways and the core exocytotic machinery, enabling inhibitory GPCR signaling to directly regulate the probability of synaptic vesicle fusion. More broadly, presynaptic inhibition plays an important role in synaptic integration by dynamically regulating neurotransmitter release probability and thereby shaping how synaptic inputs are filtered and transmitted through neural circuits.^42^

The combination of our computational predictions with the experimentally determined cryo-EM map provides new hypotheses that can be tested through various experimental approaches. For example, our model suggests the N-termini of Gβγ occupy the portion of synaptobrevin that is missing from the partially zipped SNARE construct. This could be tested by evaluating the ability of the synaptobrevin peptide corresponding to the truncation (residues 67 - 89) to compete with full-length Gβ1γ2 for binding to partially zipped SNARE. Competition would indicate that the C-terminal SNARE domain of synaptobrevin displaces the N-termini of Gβγ from a shared binding site at the C-terminus of the SNARE complex, in support of our current model.

## CONCLUSION

On the basis of our structural and biochemical studies, we propose a molecular mechanism by which Gβ1γ2 disrupts N-to-C-terminal zippering of the SNARE complex by impeding synaptobrevin incorporation into the core SNARE helical bundle. Gβγ may alter the energetic favorability of SNARE complex formation by either stabilizing the disordered C-terminus of SNAP-25 or by triggering a disorder-to-order transition. This could reduce the free-energy released by synaptobrevin incorporation, which orders the C-terminus of the SNARE complex into a highly stable four helical bundle.^23,26,43,44^ Moreover, Gβ1γ2 also disrupts synaptotagmin binding to the SNARE complex, although it is not yet clear whether this occurs via allostery or competition for a shared binding site.^3,6^ While our structure did not reach high-resolution, we are still able to yield essential mechanistic insight.

